# Topographic CA1 input shapes subicular spatial coding

**DOI:** 10.64898/2026.03.24.714092

**Authors:** Yanjun Sun, Daniel T. Pederick, Xiangmin Xu, Liqun Luo, Lisa M. Giocomo

**Author notes:** Corresponding authors (LL), (LMG).

## Abstract

Precise anatomical topographic mapping in the brain supports high-fidelity information transfer for computations such as sensory alignment and motor coordination^1,2^. As a brain structure central to learning, memory, and navigation, the hippocampal formation is likewise organized with striking topographic precision^3–8^. Within this circuit, the subiculum plays a critical role in forming the cognitive map by integrating spatial and non-spatial information from both CA1 and the entorhinal cortex (EC)^9–17^. Inputs from these regions follow a continuous topography such that distal subiculum receives projections from proximal CA1 (pCA1) and medial EC (MEC), whereas proximal subiculum receives projections from distal CA1 (dCA1) and lateral EC (LEC)^3–5,7,8^. This anatomical architecture suggests that subicular computations may depend critically on the fidelity of these circuit topographies. However, because CA1 and EC provide partially overlapping spatial information, it remains unclear to what extent CA1-to-subiculum topography, independent of EC’s contribution, shapes the physiological properties of subicular neurons. Here, by selectively disrupting CA1-to-subiculum topographic projections in *latrophilin-2* conditional knockout mice, we show that precise topography shapes the anatomical distribution of subicular spatial coding while preserving single-cell tuning. Disrupted CA1 input topography also selectively impairs subicular boundary vector coding and long-term network stability. Thus, CA1 input topography is critical for organizing subicular spatial coding and network dynamics.

**Highlights:**

- CA1→subiculum topography organizes spatial coding distribution in the subiculum.
- Disrupted CA1 input topography spares individual subicular place cell tuning.
- Disrupted CA1 input topography selectively impairs subicular boundary vector coding.
- Long-term subicular ensemble reactivation requires precise CA1 input topography.

## Results

A central challenge in dissecting subicular function is that convergent and divergent pathways are embedded within a densely interconnected network^6,18^. The discovery of *teneurin-3 (Ten3)*–*latrophilin-2 (Lphn2)* – mediated wiring rules governing CA1→subiculum topography has made it possible to selectively perturb this circuit organization^19,20^. Here, we conditionally knocked out *Lphn2* in subicular excitatory neurons by crossing floxed *Lphn2* transgenic mice (*Lphn2^fl/fl^*)^21^ to a *Nts-Cre* line^22^, in which Cre recombinase expression is primarily restricted to subicular excitatory neurons^23^. As shown in previous studies, this manipulation selectively disrupts the topographic organization of CA1→subiculum projection: instead of targeting precisely to distal subiculum in wild-type animals, axons from proximal CA1 neurons also spread to proximal subiculum (Figure 1A)^20^. In contrast, the precision of EC → subiculum projection shows no detectable change (Figure 1A)^23^. This allows us to test the functional role of CA1→subiculum topography.

**Figure 1.**
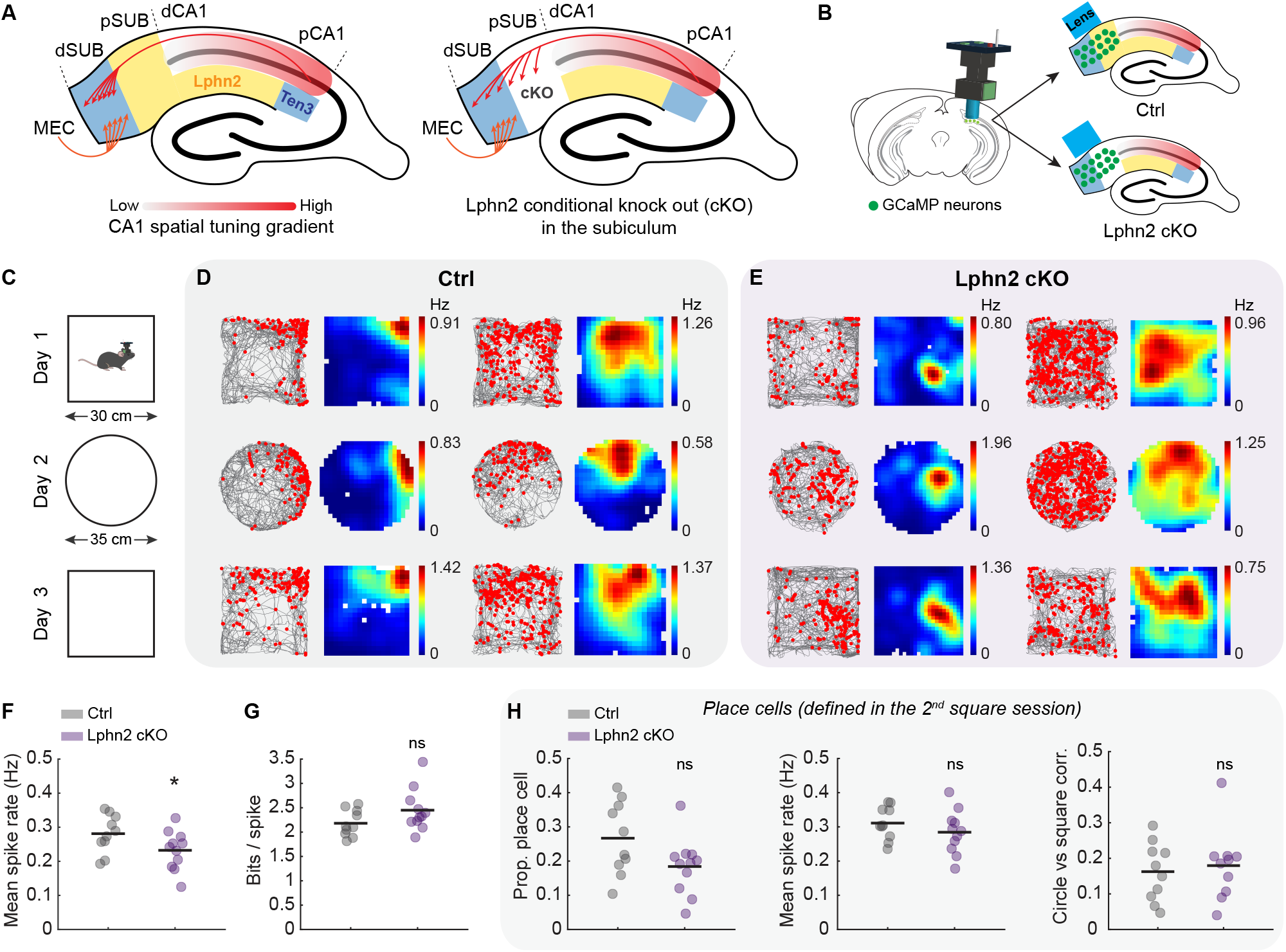
Subicular place cells preserved their spatial tuning in *Lphn2* cKO mice. **(A)** Schematic of CA1-to-subiculum projection topography in wild-type and *Lphn2* cKO mice. In wild-type mice (left), proximal CA1 axons project preferentially to distal subiculum. In subiculum-specific *Lphn2* conditional knockout mice (right), proximal CA1 axons ectopically spread into proximal subiculum. Blue indicates *Ten3* expression, yellow indicates *Lphn2* expression, and arrows represent axonal projections. **(B)** Schematic of miniscope calcium imaging in the subiculum for Ctrl and *Lphn2* cKO mice. **(C)** Open arena environment shapes. **(D)** Two representative place cells from two control (Ctrl) mice. Each column represents a single place cell, with its activity tracked across multiple sessions. The raster plot (left) shows deconvolved spikes (red dots) overlaid on the animal’s running trajectory (grey lines), while the spatial rate map (right) is color-coded, with red indicating the maximum and blue indicating the minimum values. **(E)** Same as D, but for *Nts-Cre;Lphn2^fl/fl^* (*Lphn2* cKO) mice. **(F)** Comparison of mean spike rate across all recorded neurons from Ctrl and *Lphn2* cKO mice. Each dot represents a mouse (Wilcoxon rank-sum test: *p =* 0.045; Ctrl: *n* = 10 mice (total 1782 neurons); *Lphn2* cKO: *n =* 11 mice (total 2749 neurons)). The black bar represents the mean. Quantification was based on day 3 square session. **(G)** Same as F, but for the information score comparison (Wilcoxon rank-sum test: *p =* 0.084). **(H)** Comparisons of place cells properties in Ctrl and *Lphn2* cKO mice (Wilcoxon rank-sum test: all *p >* 0.05).

To record the activity of subicular neurons in freely behaving mice, we injected GCaMP6f-expressing AAV viruses into the dorsal subiculum of either *Nts-Cre;Lphn2^fl/fl^* (*Lphn2* cKO, hereafter) or *Nts-Cre;Lphn2^+/+^*(littermate control; Ctrl) mice, and performed in vivo calcium imaging using a single photon (1P) miniscope (Figure 1B; Figure S1A; STAR Methods). Ca^2+^ signals were extracted with CNMF and OASIS deconvolution, and subsequently binarized to estimate spikes for all cells (STAR Methods, Sun et al., 2022, 2024). We treated the deconvolved spikes as equivalent to electrophysiological spikes for downstream analyses.

### Subicular place cells preserved tuning but shifted proximally in *Lphn2* cKO mice

We imaged subicular neurons as mice freely explored a sequence of open arena environments – a square, a circle, and a square again – over three consecutive days (Figure 1C). Subicular neurons in Ctrl and *Lphn2* cKO mice included both sharply and broadly tuned place cells (Figures 1D and 1E). To minimize novelty effects and ensure stable spatial representations, we analyzed neural tuning properties using activity from the second square session on day 3 (except for measurements requiring across-session calculations; see below). Compared to Ctrl mice, subicular neurons (i.e., all recorded cells) in *Lphn2* cKO mice showed a slightly reduced mean spike rate (Ctrl vs. *Lphn2* cKO, mean ± standard error of the mean [SEM]: 0.28 ± 0.02 vs. 0.23 ± 0.02), but comparable spatial information (2.18 ± 0.09 vs. 2.45 ± 0.13) (Figures 1F and 1G). To capture the diverse spatial firing patterns observed in the subiculum^9,11,24–27^, we defined subicular place cells as neurons that showed significant spatial information relative to shuffled data and met a minimum mean spike-rate threshold (STAR Methods). The proportion of place cells was numerically lower in *Lphn2* cKO mice than in Ctrl mice, but this difference did not reach statistical significance (Ctrl = 0.27 ± 0.03 vs. *Lphn2* cKO = 0.18 ± 0.02) (Figure 1H). In addition, key tuning properties—mean spike rate, peak spike rate, spatial information, field size, coherence, stability, and degree of ‘remapping’ (i.e., correlation between tuning curves) across the circle and square — were not significantly different (Figure 1H; Figure S1B). To further examine place cell remapping, we used a shuttle box containing two connected square boxes with different visual and tactile cues (Figures S1C and S1D). Consistent with the circle and square arena results, Ctrl and *Lphn2* cKO place cells exhibited comparable remapping across the two contexts (Figure S1E).

We next examined the anatomical distribution of place cells under the imaging window (Figure 2A). For each mouse, we first aligned miniscope images to the midline to ensure consistent proximodistal and rostrocaudal orientation across mice (Figure 2A; Figure S1F). To calculate place cell density, we identified the anatomical location of each place cell and normalized the number of place cells to the total number of recorded neurons in each image bin (Figures 2B–2E). Thus, place cell density ranged from 0 to 1 (Figures 2C and 2E), with 1 representing the maximum possible density. For each mouse, the final place cell density was obtained by averaging the densities across the square, circle, and shuttle box conditions (Figure 2F). We then compared this final density distribution along either the proximodistal or rostrocaudal axis across different mice (Figures 2G–2K). Compared to Ctrl mice, *Lphn2* cKO mice showed a significant shift in the anatomical distribution of place cells toward the proximal subiculum (peak bin location, mean ± SEM: Ctrl vs. *Lphn2* cKO: 11.30 ± 0.65 vs. 7.36 ± 1.04) (Figures 2G–2I), with no changes observed along the rostrocaudal axis (8.90 ± 0.48 vs. 8.36 ± 0.92) (Figures 2J and 2K). To confirm this finding, we repeated the same analysis within each environment (without averaging across conditions) and observed a consistent proximal shift of place cells in all environments (Figures S2A–S2I). Furthermore, an alternative method of defining place cells also yielded similar results (Figures S2J–S2L; STAR Methods).

**Figure 2.**
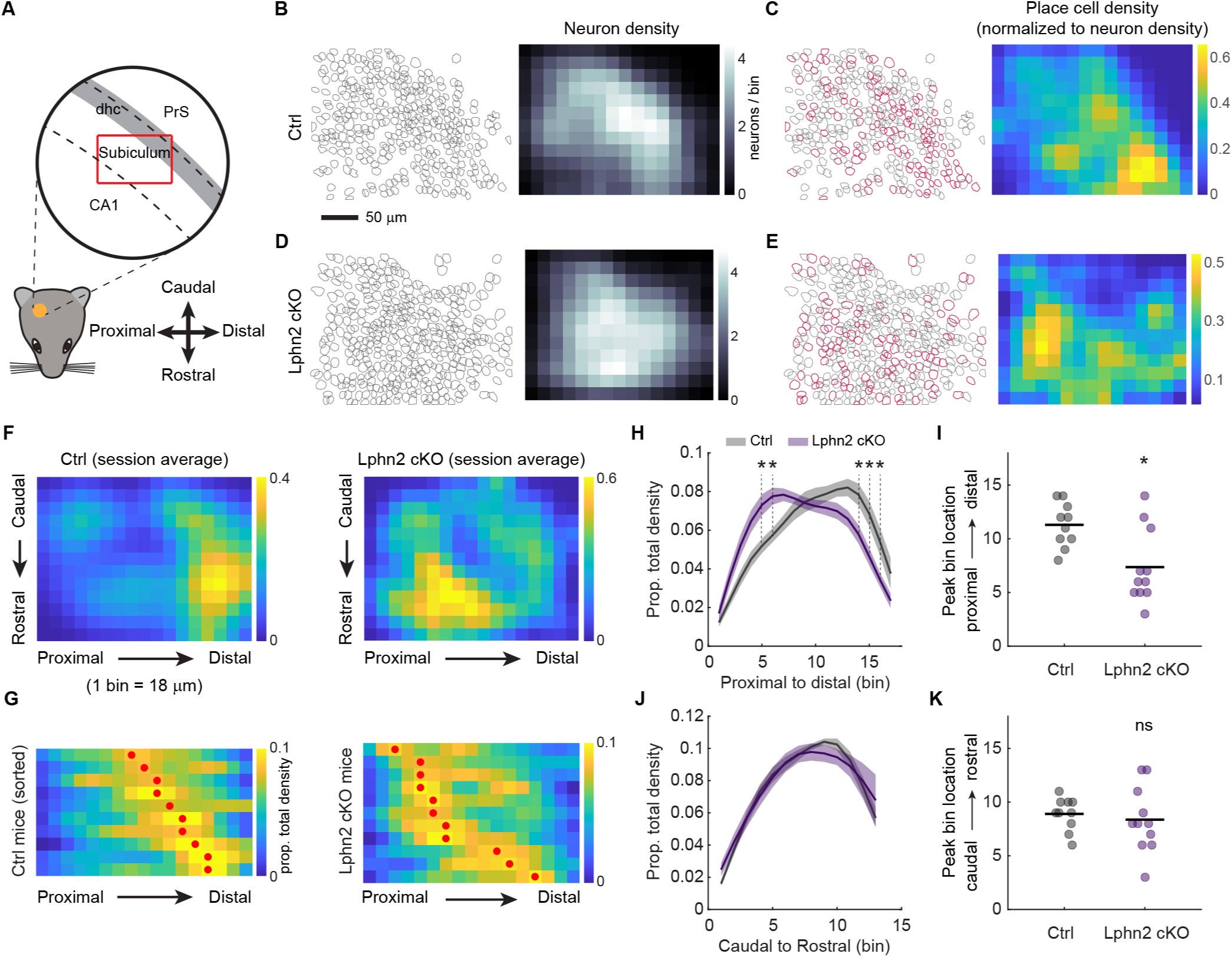
Subicular place cells shifted anatomically to the proximal subiculum. **(A)** Illustration of the anatomical structure under the miniscope imaging window. The circle represents the implanted GRIN lens (1.8 mm diameter). The red box represents the field of view of the miniscope CMOS sensor, which is 700 ìm × 450 μm. PrS: presubiculum; dhc: dorsal hippocampal commissure. **(B)** Left: anatomical footprints of all recorded subicular neurons from a representative Ctrl mouse in a single session. Each grey contour represents a neuron. Right: corresponding density map, with neuron density ranging from low (black) to high (white). Spatial bin size: 18 × 18 μm. **(C)** Left: distribution of identified place cells (red) in the subiculum from the same mouse as in I. Right: density map of place cells, normalized by the density of all neurons. **(D)** Same as B, but for an *Lphn2* cKO mouse. **(E)** Same as C, but for an *Lphn2* cKO mouse. **(F)** Left: place cell density map, averaged across all recording sessions (square, circle, and shuttle box), from a representative Ctrl mouse. Right: same as left, but for a representative Lphn2 cKO mouse. **(G)** Distribution of place cell density along the proximodistal axis for all Ctrl (left) and Lphn2 cKO (right) mice. Each row represents a mouse, with a red dot indicating the peak of its distribution. **(H)** Comparison of place cell density distribution along the proximodistal axis between Ctrl and *Lphn2* cKO mice (two-way ANOVA with Sidak’s multiple comparisons: * p-values range from 0.029 to 0.045). **(I)** Peak location of place cell density distributions (Wilcoxon rank-sum test: *p =* 0.013). **(J)** Same as H, but along the caudal-to-rostral axis (two-way ANOVA with Sidak’s multiple comparisons: all *p >* 0.05). **(K)** Same as I, but along the caudal-to-rostral axis (Wilcoxon rank-sum test: *p =* 0.46).

### *Lphn2* cKO selectively impairs subicular boundary coding while sparing corner coding

To further investigate the role of *Lphn2* in spatial coding, we examined the tuning properties of boundary vector cells (BVCs) and corner cells^9,11^ – two neuronal populations that encode geometric features of the environment.

We observed BVCs in both Ctrl and *Lphn2* cKO mice (Figures 3A and 3B). However, compared to Ctrl, BVCs in *Lphn2* cKO mice were fewer in number (Ctrl vs. *Lphn2* cKO: 0.035 ± 0.008 vs. 0.017 ± 0.004, mean ± s.e.m.), exhibited lower mean spike rates (0.32 ± 0.02 vs. 0.23 ± 0.02) but higher spatial information (2.12 ± 0.05 vs. 2.49 ± 0.14) (Figure 3C; Figure S3A). Other tuning properties of BVCs remained unchanged in the *Lphn2* cKO mice, except for a decreasing trend in the across session stability (Figure S3A). To better account for individual animal variability, we compared the tuning properties of BVCs to those of place cells within the same animal. In Ctrl mice, BVCs were more stable than place cells within- and across-sessions (place cells vs. BVCs: 0.32 ± 0.05 vs. 0.41 ± 0.05) in the square environment (Figure 3D; Figure S3B), but BVCs and place cells exhibited a similar degree of remapping between square and circle environments (place cells vs. BVCs: 0.16 ± 0.03 vs. 0.12 ± 0.04) (Figure 3D). In contrast, this relationship was reversed in *Lphn2* cKO mice, with comparable stability between BVCs and place cells in the square environment (place cells vs. BVCs: 0.23 ± 0.02 vs. 0.25 ± 0.04) (Figure 3E; Figure S3C), but stronger square to circle remapping in BVCs (place cells vs. BVCs: 0.18 ± 0.03 vs. 0.06 ± 0.03) (Figure 3E). This within-subject comparison further confirmed the decreased mean spike rate and increased spatial information of BVCs, relative to place cells, in *Lphn2* cKO mice (Figure S3C). Despite changes in BVC tuning properties, the anatomical distribution of BVCs in *Lphn2* cKO mice were not different from those in Ctrl mice (Figures 3F–3J).

**Figure 3.**
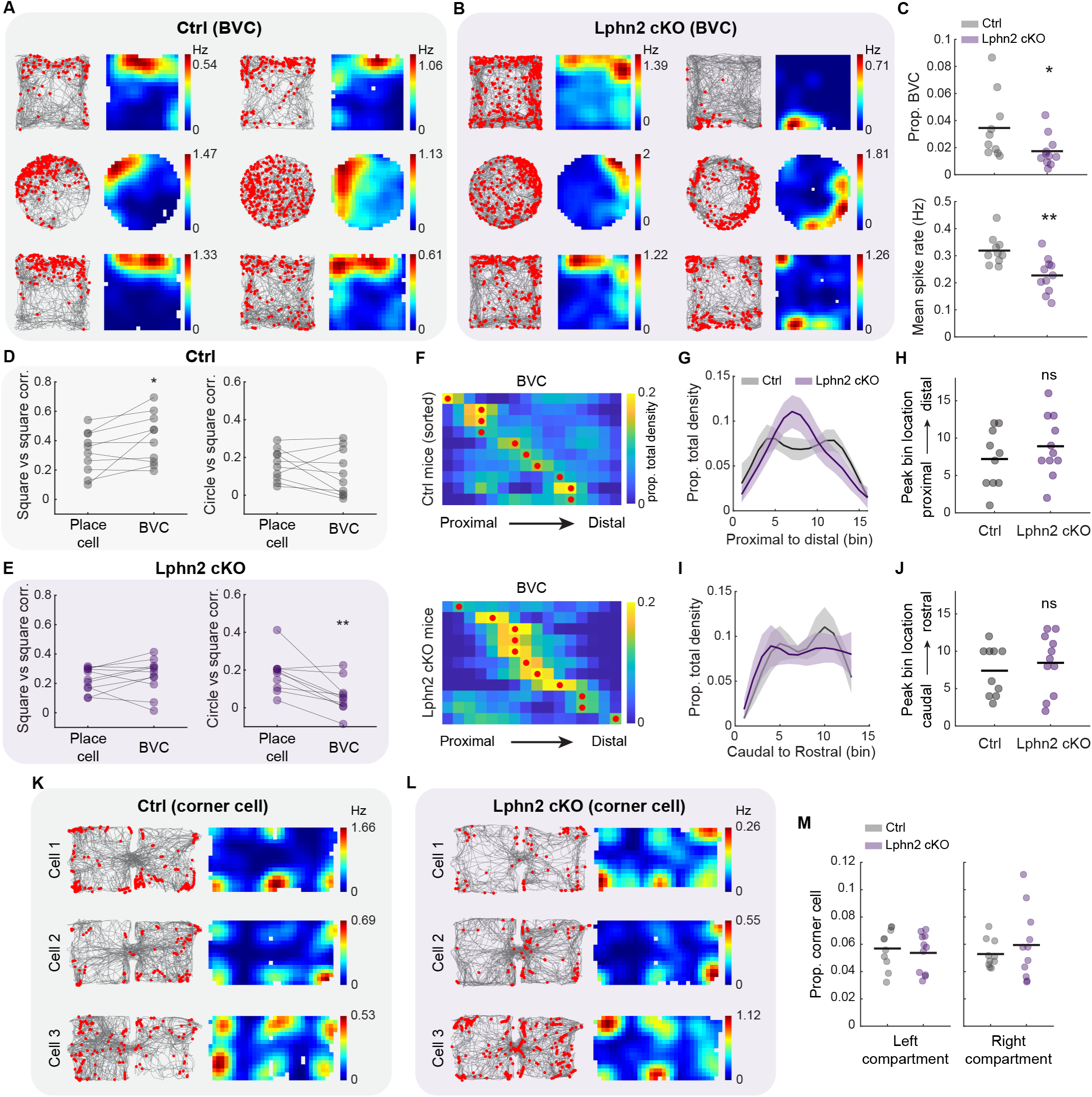
*Lphn2* cKO selectively alters subicular boundary coding while sparing corner coding. **(A)** Two representative boundary vector cells (BVCs) from two Ctrl mice, plotted as in Figure 1D. **(B)** Same as A, but for *Lphn2* cKO mice. **(C)** Top: Proportion of BVCs in Ctrl and *Lphn2* cKO mice (Wilcoxon rank-sum test: *p =* 0.022; Ctrl: *n =* 10 mice; *Lphn2* cKO: *n =* 11 mice). Bottom: Mean spike rate of BVCs from Ctrl and *Lphn2* cKO mice. Each dot represents a mouse, averaged across all square sessions (Wilcoxon rank-sum test: *p =* 0.0043). Black bar represents the mean. **(D)** Property comparisons between place cells and BVCs within Ctrl mice. Left: Rate map correlation between two square sessions (i.e., across-session stability, Wilcoxon sign-rank test: *p =* 0.027; *n =* 10 mice). Right: Rate map correlation between square and circle sessions (i.e., degree of remapping, Wilcoxon sign-rank test: *p =* 0.23). **(E)** Same as D, but for *Lphn2* cKO mice. Left, Wilcoxon sign-rank test: *p =* 0.41; *n =* 11 mice. Right, Wilcoxon sign-rank test: *p =* 0.0039; *n =* 10 mice (*note:* one mouse does not have a circle session). **(F)** Distribution of BVC density along the proximodistal axis for all Ctrl (top) and *Lphn2* cKO (bottom) mice. Each row represents a mouse, with the red dot indicating the peak of its distribution. **(G)** Comparison of BVC density distribution along the proximodistal axis between Ctrl and *Lphn2* cKO mice (two-way ANOVA with Sidak’s multiple comparisons: all *p >* 0.05). **(H)** Peak location of BVC density distributions (Wilcoxon rank-sum test: *p =* 0.38). **(I)** Same as G, but along the caudal-to-rostral axis (two-way ANOVA with Sidak’s multiple comparisons: all *p >* 0.05). **(J)** Same as H, but along the caudal-to-rostral axis (Wilcoxon rank-sum test: *p =* 0.59). **(K)** Three representative corner cells from three Ctrl mice in the shuttle box session, plotted as in Figure 1C. **(L)** Same as K, but for *Lphn2* cKO mice. **(M)** Proportion of corner cells in the left (Wilcoxon rank-sum test: *p =* 0.65) or right (Wilcoxon rank-sum test: *p =* 0.81) shuttle box compartment in Ctrl and *Lphn2* cKO mice.

Next, we identified subicular corner cells in a shuttle box environment and observed corner coding in both Ctrl and *Lphn2* cKO mice (Figures 3K and 3L). In each shuttle box compartment, the proportion of corner cells was comparable between Ctrl and *Lphn2* cKO mice (Figure 3M). Similar to BVCs, the anatomical distribution of corner cells in the subiculum remained unchanged in *Lphn2* cKO mice (Figure S3D). Together, these results suggest that subicular boundary coding is preferentially sensitive to CA1→subiculum topography.

### Head direction tuning remains unaffected in *Lphn2* cKO mice

In addition to spatial coding, subicular neurons exhibit tuning to head direction^10,26,27^. We identified head direction cells in both the Ctrl and *Lphn2* cKO mice (Figures S4A and S4B), with 60% lacking spatial modulation (i.e., they were not place cells), indicating a substantial segregation between the head direction and place cell populations. Similar to place cells, the tuning properties of head direction cells were comparable between Ctrl and *Lphn2* cKO mice (Figure S4C). However, unlike place cells, head direction cells did not show any shift in anatomical distribution along either the proximodistal or rostrocaudal axis (Figures S4D–S4I). Together, these results suggest that the CA1→subiculum circuit topography specifically contributes to the anatomical distribution of spatial, but not directional, coding in the subiculum.

### Long-term reactivation of coordinated subicular activity is reduced in *Lphn2* cKO mice

We next examined whether the population dynamics of temporally coordinated subicular cell ensembles differed between Ctrl and *Lphn2* cKO mice. We applied a PCA/ICA-based cell ensemble detection method^28^ to identify groups of neurons exhibiting temporally coordinated activity (Figure 4A; STAR Methods), and compared the ensemble properties between Ctrl and *Lphn2* cKO mice while controlling the sampling variability (i.e., subsampled, STAR Methods). As expected, neurons assigned to coordinated ensembles exhibited stronger pairwise correlations than the full neuronal population in both Ctrl and *Lphn2* cKO mice (Figures S5A and S5B). On average, nearly 50% of the subsampled neurons were grouped into approximately a dozen distinct ensembles (Figure S5). Basic ensemble properties, including ensemble count, neurons per ensemble, and the number of neurons participating in more than one ensemble, were similar between genotypes (Figure S5).

**Figure 4.**
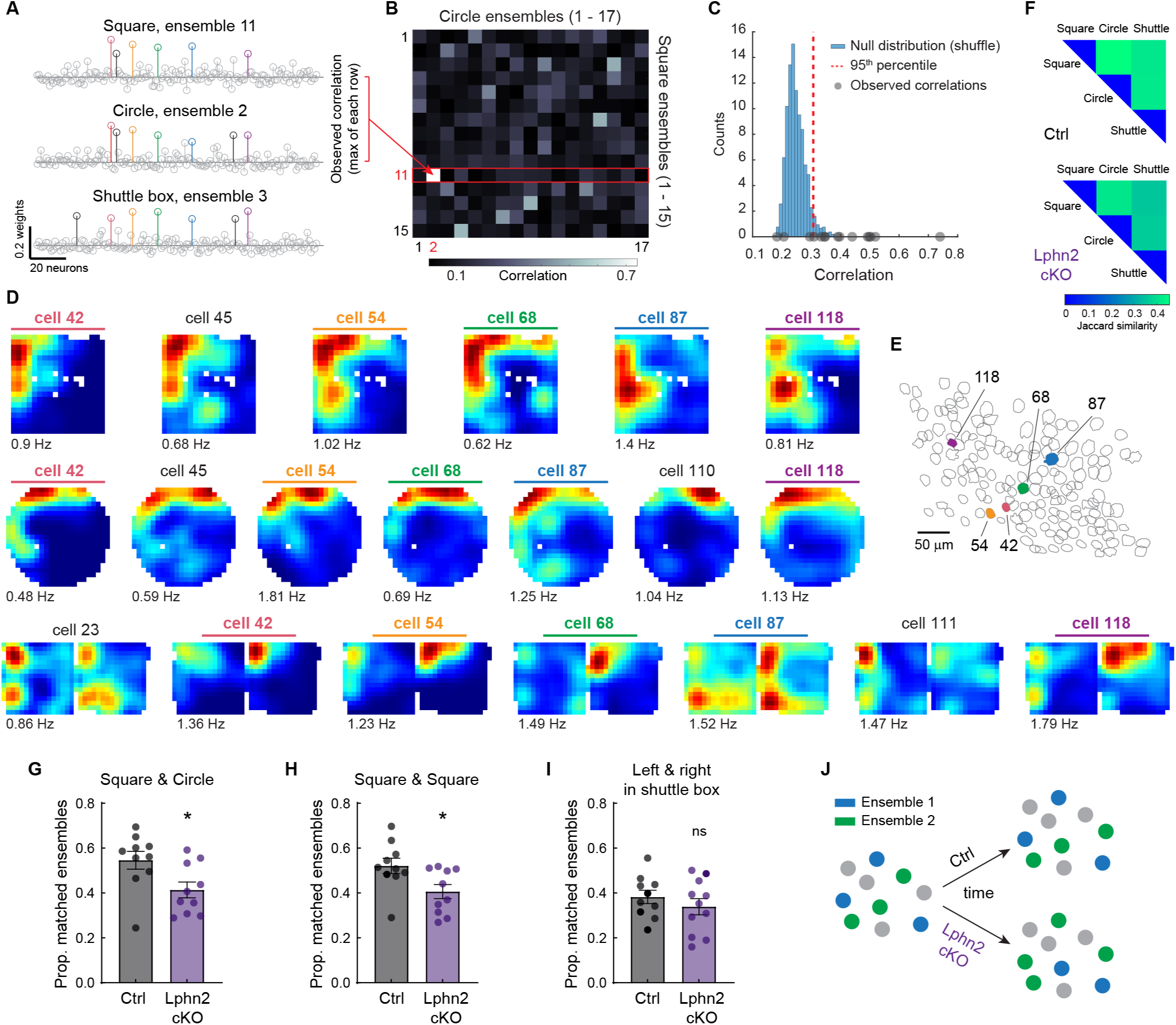
Long-term reactivation of coordinated subicular activity is reduced in *Lphn2* cKO mice. **(A)** Vectors of ensemble weights for a representative reoccurring ensemble across three recording sessions (square, circle, and shuttle box). In each vector, non-gray lollipop bars indicate neurons that significantly contribute to the ensemble, while colored bars highlight neurons consistently present in this ensemble across all three environments. **(B)** Correlation matrix of ensemble weights between a square and a circle session from a representative mouse, color-coded from high (white) to low (black). The highest correlation value for each row was used to identify potential matches between ensembles across sessions (see observed correlations in C). The example entry pointed by the red arrow represents the correlation between the first two ensemble weights in A. **(C)** Criteria for matching ensembles across sessions using shuffled correlation values. Two ensembles, from different recording sessions, were considered the same if their observed correlation (gray dots) exceeded the 95^th^ percentile of the shuffled distribution. **(D)** Spatial rate maps of the ensemble neurons shown in A, with neuron numbers color-coded as in A. **(E)** Anatomical locations of neurons that are consistently present in the representative ensemble shown in A. **(F)** Overlap of neurons classified into neural ensembles across different environments (top: Ctrl; bottom: *Lphn2* cKO). The heatmap is color-coded from high (green) to low Jaccard similarity (blue). Higher similarity indicates that the same neurons are consistently assigned to neural ensembles across environments, regardless of which specific ensemble they belong to. Note that diagonal entries are set to zero for better visualization. Ctrl vs. *Lphn2* cKO, square & circle: 0.46 ± 0.03 vs. 0.41 ± 0.02, mean ± s.e.m.; square & shuttle box: 0.41 ± 0.03 vs. 0.35 ± 0.03; circle & shuttle box: 0.41 ± 0.02 vs. 0.36 ± 0.02 (Wilcoxon rank-sum test: all *p >* 0.05). **(G)** Proportion of matched ensembles in Ctrl and *Lphn2* cKO mice between a square and a circle session (Wilcoxon rank-sum test: *p =* 0.018; Ctrl: *n =* 10 mice, *Lphn2* cKO: *n =* 10 mice). **(H)** Same as G, but between two square sessions (Wilcoxon rank-sum test: *p =* 0.043). **(I)** Proportion of matched ensembles in Ctrl and *Lphn2* cKO mice between the left and right compartments in the shuttle box (Wilcoxon rank-sum test: *p =* 0.47; Ctrl: *n =* 10 mice, *Lphn2* cKO: *n =* 11 mice). The left and right compartments were connected, allowing the mouse to freely explore both compartments in a single session. **(J)** Schematic illustrating the more pronounced reconfiguration of ensemble assignments in *Lphn2* cKO mice compared to Ctrl.

To assess how coordinated ensembles evolve over time in the same animal, we tracked whether the same ensembles reappeared across different sessions (Figures 4A–4C; Methods). To identify the same ensemble across sessions, we computed, for a given ensemble, the correlation between its weight vector and those of all ensembles in another session, identifying the most strongly correlated pair (Figures 4A and 4B). If the observed correlation of the pair exceeded the 95^th^ percentile of a shuffled distribution, we considered them to be the same ensemble (Figure 4C; STAR Methods). An example is shown in Figure 4A and 4D, where three ensembles recorded from different sessions of a single mouse were successfully matched. Despite being anatomically distant from each other, most neurons maintained their membership in this ensemble across sessions, with only a few joining or dropping out over time (Figures 4A, 4D and 4E). We refer to this phenomenon—when an ensemble from one session can be matched to one from another session in the same animal—as ensemble reoccurrence, which reflects the stability of the participating neuron group over time.

To distinguish this from general neuron turnover, we examined the overlap of neurons that participated in any ensemble across sessions. This metric captures how many of the same neurons were recruited into any ensemble across different environments (Figure 4F). *Lphn2* cKO mice showed a similar degree of overlap in ensemble-participating neurons across environments compared to Ctrl mice, indicating that a comparable set of neurons was recruited into ensembles with a similar turnover rate across environments in both genotypes (Figure 4F). However, when we examined ensemble reoccurrence across sessions, a significant reduction of matched ensembles was observed in *Lphn2* cKO mice (Figure 4G). This decrease was seen in both square-circle (Ctrl vs. *Lphn2* cKO: 0.55 ± 0.04 vs. 0.41 ± 0.04, mean ± s.e.m.) (Figure 4G) and square-square (0.52 ± 0.03 vs. 0.41 ± 0.03) (Figure 4H) comparisons, suggesting that the disruption is not context-dependent but rather reflects an internal deficit in maintaining long-term ensemble structure. Importantly, this reduction in ensemble reoccurrence was not observed when comparing recordings from the connected left and right compartments of a shuttle box (Ctrl vs. *Lphn2* cKO: 0.38 ± 0.03 vs. 0.34 ± 0.04) (Figure 4I). This suggests that the disruption in ensemble reoccurrence emerges over extended timescales (i.e., hours to days), rather than being immediately apparent within a single recording session (Figure 4J). Together, these findings demonstrate that proper CA1→subiculum topography is required for sustaining long-term temporal coordination in the subicular neural network.

## Discussion

Neurons in pCA1 and MEC, including place cells and grid cells, provide highly spatially tuned inputs that converge preferentially onto distal subiculum^12,14^, whereas dCA1 and LEC preferentially target proximal subiculum and are associated with relatively greater non-spatial coding^14,16,17^. This organization suggests that the proximodistal axis of CA1–subiculum connectivity may help distribute distinct computations across the subiculum. Although pCA1 and MEC both carry spatial information and could, in principle, compensate for one another, our results show that the precise topography of CA1 projections directly constrains the anatomical distribution of spatial coding in the subiculum, while leaving the tuning properties of individual place cells largely preserved. Thus, the CA1→subiculum projection topography appears to organize where spatially tuned subicular neurons are expressed along the proximodistal axis, and MEC inputs alone are insufficient to maintain this distribution. This topographic organization may be particularly important for specialized spatial computations, including boundary vector coding and the long-term network stability of subicular population activity. In contrast, the preservation of subicular head direction coding indicates that directional information operates via a distinct, CA1-independent pathway. Collectively, our results reveal that the topographic organization of CA1 inputs is not simply an anatomical feature, but a functional organizing principle for distributing and stabilizing spatial representations in the subiculum.

Although inputs from MEC to the subiculum and pCA1 were intact in *Lphn2* cKO mice^23^, we cannot fully exclude the possibility of indirect effects that could arise from changes in other subicular circuits. A known circuit alteration in this mouse line is the mistargeting of projections from the subiculum to the medial mammillary nucleus (MMN)^23^. Because MMN projects to the anterior thalamic nuclei (ATN), which broadcast head direction signals to cortical regions including the subiculum^29^, such miswiring could in principle alter subicular coding. However, we found that subicular head direction coding in *Lphn2* cKO mice was unaffected, with tuning properties and anatomical distribution comparable to controls. Thus, disruption of the subiculum → MMN pathway is unlikely to explain our findings. These results also suggest that ATN → subiculum projections remain functionally preserved under this manipulation^30^. In addition to their roles in topographic wiring, *teneurins* and *latrophilins* have been implicated in synapse formation, including a role for *Lphn2* in EC → CA1 synapse assembly^21,31^. Although synapse formation defects could in principle contribute to the phenotypes observed here, the preservation of general coding properties of spatial and head direction tuning (Figures 1F–1H) argues against a global synaptic deficit. Instead, the selective proximal shift of spatial coding is most consistent with disrupted CA1→subiculum topography.

Our results further showed that BVCs, but not corner cells, were selectively affected by loss of CA1 topography. In controls, boundary coding was more stable than place coding, consistent with a role in anchoring spatial maps across environments. This advantage was reduced in *Lphn2* cKO mice, highlighting a selective contribution of CA1 inputs to stabilizing boundary-based reference frames. One potential explanation for the reduced number of BVCs is that boundary coding requires the integration of both spatial and directional information. Possible sources of head direction input to subiculum include MEC and ATN^32–34^, both of which project preferentially to distal subiculum^8,35^. Because these directional inputs converge with spatial inputs from proximal CA1 most prominently in the distal subiculum, this region may serve as a primary site for spatial–directional integration. By shifting spatial coding proximally, disruption of CA1 topography likely reduces this integration, thereby weakening BVC emergence across the subiculum. Thus, our results suggest that subicular boundary coding requires properly organized CA1 input, rather than being directly inherited from the MEC.

At the network level, coordinated cell ensembles in *Lphn2* cKO mice formed normally but showed reduced reoccurrence across days, indicating a deficit in long-term temporal coordination. This suggests that precisely organized CA1 inputs support the reactivation of consistent subicular ensemble patterns over time, a process essential for long-term memory^36^. However, the behavioral consequences of disrupting CA1→subiculum topography remain to be determined. Together, these findings demonstrate that *Ten3*–*Lphn2*–mediated wiring rules^23^ not only establish topographic maps but also shape hippocampal computations and dynamics.

## STAR METHODS

### KEY RESOURCES TABLE

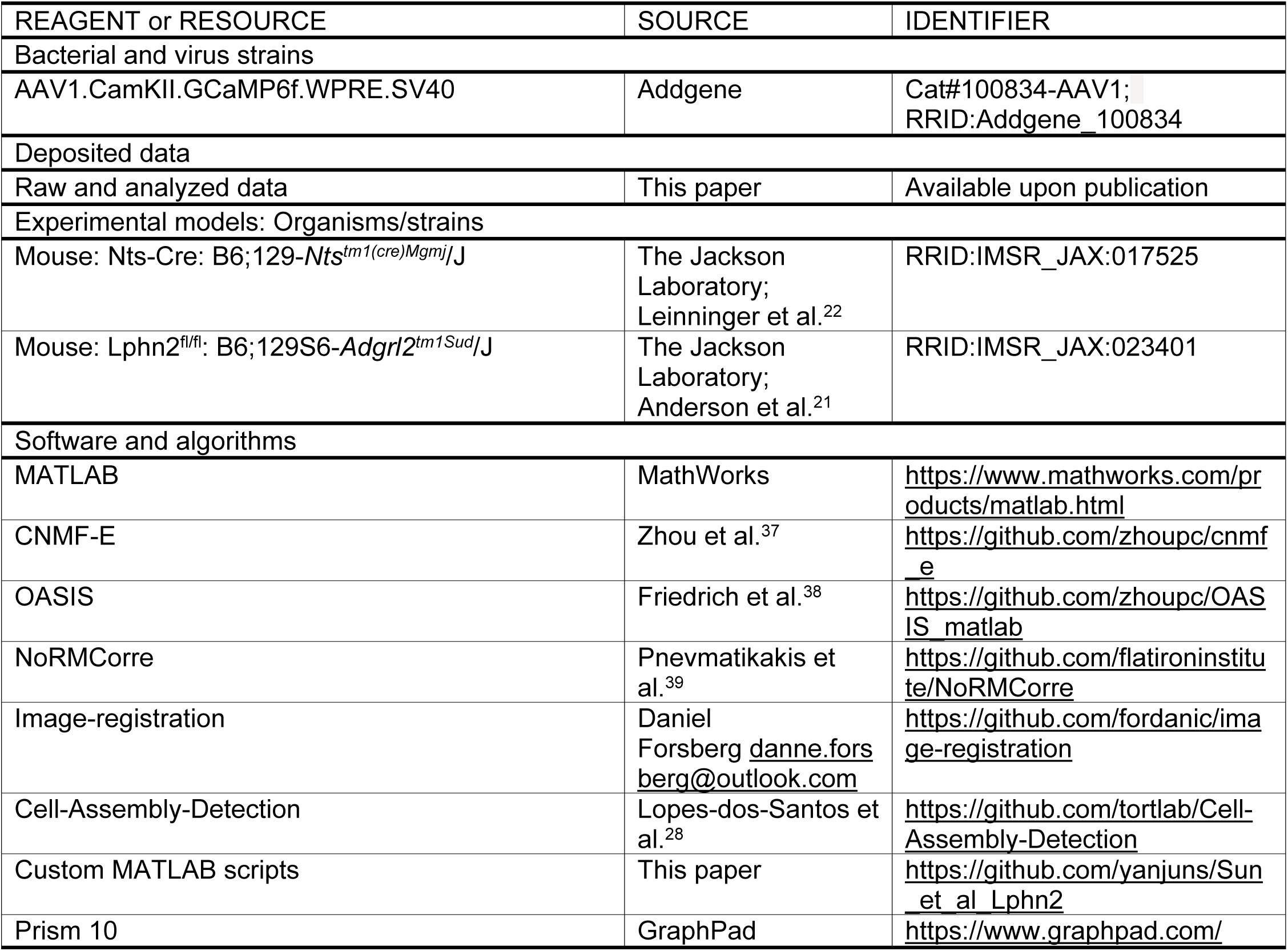

### EXPERIMENTAL MODEL DETAILS

#### Subjects

All procedures were conducted according to the National Institutes of Health guidelines for animal care and use and approved by the Institutional Animal Care and Use Committee at Stanford University School of Medicine and the University of California, Irvine. *Nts-Cre* mice (JAX #017525)^22^ were crossed with *Lphn2^fl/fl^*mice (JAX #023401)^21^ to generate experimental and control animals. To minimize background effects, only animals on a C57BL/6 background were used for experiments. A total of 11 *Nts-Cre;Lphn2 ^fl/fl^* (*Lphn2* cKO; 3 male and 8 female) and 10 *Nts-Cre;Lphn2^+/+^*(littermate control; 6 male and 4 female) mice were used for subiculum Ca^2+^ imaging. Mice were group housed with same-sex littermates until the time of surgery. After surgery mice were singly housed at 21-22 °C and 29-41% humidity. Mice were kept on a 12-hour light/dark cycle and had ad libitum access to food and water in their home cages at all times. All experiments were carried out during the light phase. Data from both males and females were combined for analysis, as we did not observe sex differences in, for example, place cell proportion and their anatomical distribution.

### METHODS DETAILS

#### Miniscope surgeries

Mice were 8-12 weeks old at the time of surgery. Mice were anesthetized with continuous 1-1.5% isoflurane and head fixed in a rodent stereotax. A three-axis digitally controlled micromanipulator guided by a digital atlas was used to determine bregma and lambda coordinates. To express GCaMP in the subiculum, 500 nl of AAV1-Camk2a-GCaMP6f (∼ 3.2 × 10^12^ genome copies per ml) was injected in the right subiculum at anteroposterior (AP): −3.40 mm; lateromedial (ML): +1.88 mm; and dorsoventral (DV): −1.70 mm. One week after the virus injection, we implant the gradient refractive index (GRIN) lens above the subiculum. A 1.8 mm-diameter circular craniotomy was made over the posterior cortex (centered at −3.28 mm anterior/posterior and +2 mm medial/lateral, relative to bregma). The dura was then gently removed and the cortex directly below the craniotomy aspirated using a 27- or 30-gauge blunt syringe needle attached to a vacuum pump under constant irrigation with sterile saline. The aspiration removed the corpus callosum and part of the dorsal hippocampal commissure above the imaging window but left the alveus intact. Excessive bleeding was controlled using a hemostatic sponge that had been torn into small pieces and soaked in sterile saline. The GRIN lens (0.25 pitch, 0.55 NA, 1.8 mm diameter and 4.31 mm in length, Edmund Optics) was then slowly lowered with a stereotaxic arm to the subiculum to a depth of −1.75 mm relative to the measurement of the skull surface at bregma. The GRIN lens was then fixed with cyanoacrylate and dental cement. Kwik-Sil (World Precision Instruments) was used to cover the lens at the end of surgery. Two weeks after the implantation of the GRIN lens, a small aluminum baseplate was cemented to the animal’s head on top of the existing dental cement. Specifically, Kwik-Sil was removed to expose the GRIN lens. A miniscope was then fitted into the baseplate and locked in position so that the GCaMP-expressing neurons and visible landmarks, such as blood vessels, were in focus in the field of view. After the installation of the baseplate, the imaging window was fixed for long-term, in respect to the miniscope used during installation. Thus, each mouse had a dedicated miniscope for all experiments. When not imaging, a plastic cap was placed in the baseplate to protect the GRIN lens from dust and dirt.

#### Behavioral experiments with imaging

After mice had fully recovered from the surgery, they were handled and allowed to habituate to wearing the head-mounted miniscope by freely exploring an open arena for 20 minutes every day for one week. The actual experiments took place in a different room from the habituation. The behavior rig, an 80/20 built compartment, in this dedicated room had two white walls and one black wall with salient decorations as distal visual cues, which were kept constant over the course of the entire study. For experiments described below, all the walls of the arenas were acrylic and were tightly wrapped with black paper by default to reduce potential reflections from the LEDs on the scope. A local visual cue was always available on one of the walls in the arena. In each experiment, the floors of the arenas were covered with corn bedding. All animals’ movements were voluntary.

##### Square – circle – square

This set of experiments was carried out in both a square and a circular environment. The square environment measured 30 × 30 cm, and the circle had a diameter of 35 cm. The height of all environments was 30 cm. For each mouse, we recorded one session (20 min / session) per day, following the order of square – circle – square across three days. One *Lphn2* cKO mouse did not have recordings from the circle environment. For each mouse, data from this set of experiments were aligned and concatenated, and the activity of neurons were tracked across the sessions. As described above, all the walls of the arenas were black. A local visual cue (strips of white masking tape) was present on one wall of each arena, covering the top half of the wall.

##### Shuttle box

The shuttle box consisted of two connected, 25 (L) x 25 (W) x 25 (H) cm compartments with distinct colors and visual cues. The opening in the middle was 6.5 cm wide, so that the mouse could easily run between the two compartments during miniscope recordings. The black compartment was wrapped in black paper, but not the grey compartment. For each mouse, we recorded one session (20 min / session) per day for two days.

#### Miniscope imaging data acquisition and pre-processing

Technical details for the custom-constructed miniscopes and general processing analyses are described in^11,40,41^ and at miniscope.org. Briefly, this head-mounted scope had a mass of about 3 grams and a single, flexible coaxial cable that carried power, control signals, and imaging data to the miniscope open-source Data Acquisition (DAQ) hardware and software. In our experiments, we used Miniscope V3, which had a 700 μm x 450 μm field of view with a resolution of 752 pixels x 480 pixels (∼1 μm per pixel). For subiculum imaging, we measured the effective image size (the area with detectable neurons) for each mouse and combined this information with histology. The anatomical region where neurons were recorded was approximately within a 450-μm diameter circular area centered around AP: −3.40 mm and ML: +2 mm. Due to the limitations of 1-photon imaging, we believe the recordings were primarily from the deep layer of the subiculum. Images were acquired at ∼30 frames per second (fps) and recorded to uncompressed avi files. The DAQ software also recorded the simultaneous behavior of the mouse through a high-definition webcam (Logitech) at ∼30 fps, with time stamps applied to both video streams for offline alignment.

For each set of experiments, miniscope videos of individual sessions were first concatenated and down-sampled by a factor of 2, then motion corrected using the NoRMCorre MATLAB package^39^. To align the videos across different sessions for each animal, we applied an automatic 2D image registration method (github.com/fordanic/image-registration) with rigid x-y translations according to the maximum intensity projection images for each session. The registered videos for each animal were then concatenated together in chronological order to generate a combined data set for extracting calcium activity.

To extract the calcium activity from the combined data set, we used extended constrained non-negative matrix factorization for endoscopic data (CNMF-E)^37,42^, which enables simultaneous denoising, deconvolving and demixing of calcium imaging data. A key feature includes modeling the large, rapidly fluctuating background, allowing good separation of single-neuron signals from background, and separation of partially overlapping neurons by taking a neuron’s spatial and temporal information into account (see^37^ for details). A deconvolution algorithm called OASIS^38^ was then applied to obtain the denoised neural activity and deconvolved spiking activity. These extracted calcium signals for the combined data set were then split back into each session according to their individual frame numbers. As the combined data set was large (> 5 GB), we used the Sherlock HPC cluster hosted by Stanford University to process the data across 8-12 cores and 600-700 GB of RAM. While processing this combined data set required significant computing resources, it enhanced our ability to track cells across sessions from different days. This process made it unnecessary to perform individual footprint alignment or cell registration across sessions. The position, head direction and speed of the animals were determined by applying a custom MATLAB script to the animal’s behavioral tracking video. Time points at which the speed of the animal was lower than 2 cm/s were identified and excluded from further analysis. We then used linear interpolation to temporally align the position data to the calcium imaging data.

#### Place cell analyses

##### Calculation of spatial rate maps

After obtained the deconvolved spiking activity of neurons, we binarized it by applying a threshold using a 3x standard deviation of all the deconvolved spiking activity for each neuron. The position data was sorted into 1.6 x 1.6 cm non-overlapping spatial bins. The spatial rate map for each neuron was constructed by dividing the total number of calcium spikes by the animal’s total occupancy in a given spatial bin. The rate maps were smoothed using a 2D convolution with a Gaussian filter that had a standard deviation of 2.

##### Spatial information and identification of place cells

To quantify the information content of a given neuron’s activity, we calculated spatial information scores in bits/spike (i.e., calcium spike) for each neuron according to the following formula^43^,

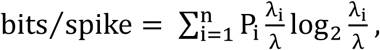

where P_i_ is the probability of the mouse occupying the i-th bin for the neuron, λ_i_ is the neuron’s unsmoothed event rate in the i-th bin, while λ is the mean rate of the neuron across the entire session. Bins with total occupancy time of less than 0.1 second were excluded from the calculation. To identify place cells, the timing of calcium spikes for each neuron was circularly shuffled 1000 times and spatial information (bits/spike) recalculated for each shuffle. This generated a distribution of shuffled information scores for each individual neuron. The value at the 95^th^ % of each shuffled distribution was used as the threshold for classifying a given neuron as a place cell, and we excluded cells with an overall mean spike rate less than the 5^th^ % of the mean spike rate distribution (i.e., ∼0.1 Hz) of all the neurons in that animal. Furthermore, we also classified place cells by adding an additional coherence criterion (coherence > 0.25) ^44–47^, and obtained consistent results (Figures S2J–S2L).

##### Neuron and place cell density and their distribution

The imaging area was divided into 18 × 18 µm spatial bins, and the centroid of each recorded neuron was determined from its CNMF-E footprint. Neuron density was calculated as the number of recorded neurons whose centroids fell within a given bin. Place cell density was defined as the proportion of these neurons classified as place cells within each bin. To generate place cell distributions along the proximodistal or rostrocaudal axis, the 2D place cell density map was rotated to align with the brain’s midline (Figure S1F). Rotation was applied only when the correction required exceeded 20°. Because the effective imaging field of view varied slightly across mice, the resulting density maps differed in size. The median for the size difference across all maps was two spatial bins. To better align distributions across animals, smaller maps were resized to match the largest map using interpolation. The 2D density map (matrix) was then collapsed into a one-dimensional vector by summing values across columns or rows for proximodistal or rostrocaudal comparisons, respectively. Each entry in the resulting vector was normalized by the total density of the map to allow comparisons across genotypes. The peak bin location within each distribution vector was also determined.

#### Head direction cell analyses

To define head direction cells, we first plotted each neuron’s firing rate as a function of the mouse’s head direction, divided into 6° bins and smoothed with a 30° mean window filter. Peak firing rate was defined as the bin with the highest rate in the polar firing rate map. The strength of directional tuning was quantified by the mean Rayleigh vector length of the tuning curve, recorded as the head direction score. Neurons were classified as head direction cells if their head direction score exceeded the 99^th^ % of the shuffled distribution and their mean spike rate was greater than 0.05 Hz. Anatomical distributions and peak bin locations were determined using the same procedure as for place cell analysis.

#### Boundary vector cell (BVC) analyses

Rate maps of all the neurons were generated using the same procedure as for place cells. To detect boundary vector cells (BVCs), we used a method based on border scores, which we calculated as described previously^48,49^:

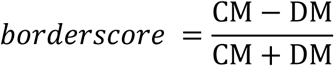

where CM is the proportion of high firing-rate bins located along one of the walls and DM is the normalized mean product of the firing rate and distance of a high firing-rate bin to the nearest wall. We identified BVCs as cells with a border score above 0.6 and whose largest field covered more than 70% of the nearest wall. Additionally, BVCs needed to have significant spatial information (i.e., as in place cells). Anatomical distributions and peak bin locations were determined using the same procedure as for place cell analysis.

#### Corner cell analyses

Corner cells were determined using previously established corner score method, as described in^11^. In brief, we first applied a threshold to filter the rate map. After filtering, each connected pixel region was considered a place field, and the x, y coordinates of the regional maxima for each field were the locations of the fields. We used a filtering threshold of 0.3 times the maximum spike rate for identifying corner cells. The coordinates of the centroid and corners of the environments were automatically detected with manual corrections. For each field, we defined the corner score as:

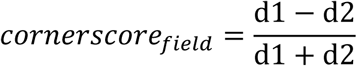

where d1 is the distance between the environmental centroid and the field, and d2 is the distance between the field and the nearest environmental corner. The score ranges from −1 for fields situated at the centroid of the arena to +1 for fields perfectly located at a corner.

##### Corner score for each cell

There were two situations that needed to be considered when calculating the corner score for each cell. First, if a cell had *n* fields in an environment that had *k* corners (*n* ≤ *k*), the corner score for that cell was defined as:

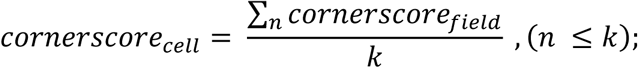

Second, if a cell had more fields than the number of environmental corners (*n* > *k*), the corner score for that cell was defined as the sum of the top *k^th^* corner scores minus the sum of the absolute values of the corner scores for the extra fields minus one, and divided by *k*. Namely,

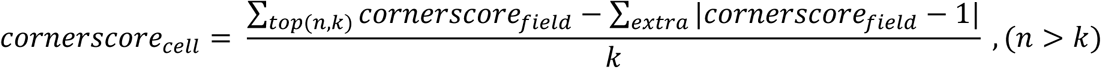

where *top(n,k)* indicates the fields (also termed ‘major fields’) that have the top *k^th^* cornerscore_field_ out of the *n* fields, and ‘extra’ refers to the corner scores for the remaining fields. In this case, the absolute values of the corner scores for the extra fields were used to penalize the final corner score for the cell, so that the score decreased if the cell had too many fields. The penalty for a given extra field ranged from 0 to 2, with 0 for the field at the corner and 2 for the field at the center. As a result, as the extra field moves away from a corner, the penalty for the overall corner score gradually increases.

##### Final definition of corner cells

To classify a corner cell, the timing of calcium spikes for each neuron was circularly shuffled 1000 times. For each shuffle, spike times were shifted randomly by 5 - 95% of the total data length, rate maps were regenerated and the corner score for each cell was recalculated. Note, for the recalculation of corner scores for the shuffled rate maps, we did not use the aforementioned penalization process. This is because shuffled rate maps often exhibited a greater number of fields than the number of corners, and thus applying the penalization lowers the 95^th^% score of the shuffled distribution (i.e. more neurons would be classified as corner cells). Thus, not using this penalization process in calculating shuffled corner scores kept the 95^th^ percentile of the shuffled distribution as high as possible for each cell to ensure a stringent selection criteria for corner cells. Finally, we defined a corner cell as a cell for whose corner score passed the 95^th^ percentile of the shuffled score, and whose distance between any two fields (major fields, if the number of fields > number of corners) was greater than half the distance between the corner and centroid of the environment.

#### Coordinated cell ensemble analyses

##### Cell ensemble detection

Calcium signals from all recorded neurons were converted to binary spike trains by thresholding each neuron’s deconvolved activity. Spike trains were then binned into 333 ms windows for each session. Coordinated cell ensembles were identified using a PCA/ICA pipeline adapted from^28^ as implemented in the Tortlab toolbox (github.com/tortlab/Cell-Assembly-Detection). The number of significant principal components was determined using the Marčenko–Pastur criterion. Independent Component Analysis was then applied to the significant subspace to obtain ensemble weight vectors. Each ensemble’s instantaneous activity was computed by projecting the binned spike data onto its weight vector. Ensemble membership was defined by each ensemble’s neuron weight distribution: neurons were assigned to ensembles if their weights exceeded mean ± 2.25 standard deviation, with sign respected. To control for sampling variability, ensemble detection was repeated 250 times using random subsamples of 100 neurons per mouse. Ensemble metrics (ensemble count, neurons per ensemble, and neuron overlap) were averaged across iterations for each animal.

##### Across-session ensemble matching

To track ensembles over time, we compared weight vectors across sessions. For each ensemble in session A, Pearson’s correlations were computed with every ensemble in session B, using the larger magnitude correlation after correcting for ICA sign flips. The maximum correlation was taken as the best match. A shuffled null distribution was built by randomly permuting neuron weights within each compared vector, recomputing the correlations, and taking the 95^th^ % as the session-pair’s match threshold. An ensemble was considered a reoccurring ensemble if its best across-session correlation exceeded this 95^th^-percentile threshold. Reoccurrence was quantified as the fraction of ensembles in session A with a significant match in session B. To distinguish reoccurrence from general neuron turnover, we also computed the overlap of ensemble-participating neurons across sessions (the proportion of neurons that participated in any ensemble in both sessions, irrespective of specific ensemble identity). The same procedure was applied to simultaneous recordings (left vs. right shuttle box compartments) and to temporally separated sessions (square–circle and square–square), allowing direct comparison of short-versus long-timescale stability.

#### Histology

After the imaging experiments were concluded, mice were deeply anesthetized with isoflurane and transcardially perfused with 10 ml of phosphate-buffered saline (PBS), followed by 30 ml of 4% paraformaldehyde-containing phosphate buffer. The brains were removed and left in 4% paraformaldehyde overnight. The next day, samples were transferred to 30% sucrose in PBS and stored in 4°C. At least 24 hours later, the brains were sectioned coronally into 30-µm-thick samples using a microtome (Leica SM2010R, Germany). All sections were counterstained with 10 μM DAPI, mounted and cover-slipped with antifade mounting media (Vectashield). Images were acquired by an automated fluorescent slide scanner (Olympus VS120-S6 slide scanner, Japan) under x10 magnification.

#### Data inclusion criteria and statistical analysis

We evaluated the imaging quality for each mouse before executing each set of experiments. No mice were excluded from the analyses as long as the experiments were executed. Analyses and statistical tests were performed using MATLAB (2022a) and GraphPad Prism 10. Data are presented as mean ± SEM. For normality checks, different test methods (D’Agostino & Pearson, Anderson-Darling, Shapiro-Wilk, and Kolmogorov-Smirnov) indicated only a portion of the data in our statistical analyses followed a Gaussian distribution. Thus, a two-tailed Wilcoxon rank-sum test and Wilcoxon signed-rank test were used for two-group unpaired and paired comparisons, respectively. For statistical comparisons between two groups across multiple conditions, two-way ANOVA with Sidak’s multiple comparisons was used. All statistical tests were conducted on a per-mouse basis. In cases where an experiment involved two or more sessions, the data were averaged across these sessions, as indicated in the corresponding text or figure legend. For example, in Figures 2F–2K, place cell density for each mouse was averaged across all recording sessions (square, circle, and shuttle box). In all experiments, the level of statistical significance was defined as p ≤ 0.05. For all the figures, * p ≤ 0.05, ** p ≤ 0.01, *** p ≤ 0.001, ns: not significant.

## Resource Availability

### Lead contact

(LMG)

### Materials Availability

Custom scripts for analyzing the data are available at https://github.com/yanjuns/Sun_et_al_Lphn2.

### Data Availability

Upon publication, data will be made available on a publicly accessible data repository site (e.g., figshare, DANDI, Mendeley).

## Acknowledgements

We thank A. Diaz and E. Velazquez for assistance with animal care and genotyping, and K. Hardcastle for coding assistance. This work was supported by National Institute of Health (NIH) Grants R01MH126904 (L.M.G.), R01MH130452 (L.M.G.), BRAIN Initiative U19NS118284 (L.M.G.), P50DA042012 (L.M.G.), the Vallee Foundation (L.M.G.), the James S. McDonnell Foundation (L.M.G.), the Simons Foundation 542987SPI (L.M.G.); NIH grant R01NS050835 (L.L.); NIH Grants R01NS104897 (X.X.), RF1 AG065675 (X.X.); NIH Grant K01DA058743 (Y.S.) and the Simons Foundation (SCGB Fellows-to-Faculty award 00007396; Y.S.). Y.S. is also a CPRIT Scholar supported by the Cancer Prevention & Research Institute of Texas (RR240082). D.T.P. is supported by the Simons Foundation (SFARI Fellows-to-Faculty Award); L.M.G. and L.L. are Investigators of the Howard Hughes Medical Institute (HHMI).

## Author contributions

Conceptualization (YS, LL, LMG), Methodology (YS, DTP, LL, XX), Investigation (YS), Visualization (YS), Funding acquisition (YS, LMG, XX, LL), Project administration (YS, LMG), Supervision (YS, LMG), Writing – original draft (YS, LMG), Writing – review and editing (YS, LMG, LL, DTP).

## Competing interests

The authors declare no competing interests.

## Notes

### Competing Interest Statement

The authors have declared no competing interest.

### Summary of Updates

We made significant changes in this revised manuscript as following: 1) We updated Figure 1A to include a schematic of normal CA1-to-subiculum projection topography, its disruption in Lphn2 conditional knockout mice, and the known proximodistal functional gradient in CA1 spatial coding. We also reorganized the figure structure to improve the presentation of the study's conceptual framework. 2) We revised the manuscript text throughout to clarify our interpretation of how CA1-to-subiculum projection topography contributes to the anatomical distribution and long-term stability of subicular spatial representations. In particular, we removed imprecise terminology such as 'scaffolding' and replaced 'cell assemblies' with 'cell ensembles' where appropriate 3) We performed additional analyses using a more stringent place-cell definition that incorporated a coherence criterion in addition to spatial information. Reanalysis with this stricter classification yielded results consistent with our original conclusions. 4) We added new statistical analyses addressing the nonsignificant numerical reduction in place-cell proportion in Lphn2 cKO mice, including outlier detection, permutation testing, Bayesian comparisons, and analyses excluding boundary vector cells (BVCs). These analyses support the interpretation that the manipulation selectively alters specific spatial coding subtypes rather than globally disrupting place coding. 5) We added new quantitative analyses of BVC anatomical distributions along the proximodistal axis, including bimodality coefficient analysis, whole-curve permutation testing, and variance comparisons. These analyses demonstrated no statistical evidence for consistent bimodality in control mice or altered spread in Lphn2 cKO mice. 6) Finally, we revised the Discussion to more clearly distinguish what is directly supported by our data from more speculative circuit interpretations, including the roles of CA1, MEC, and ATN inputs in supporting boundary vector coding.

