## Supplemental Figures for "Topographic CA1 input shapes subicular spatial coding"

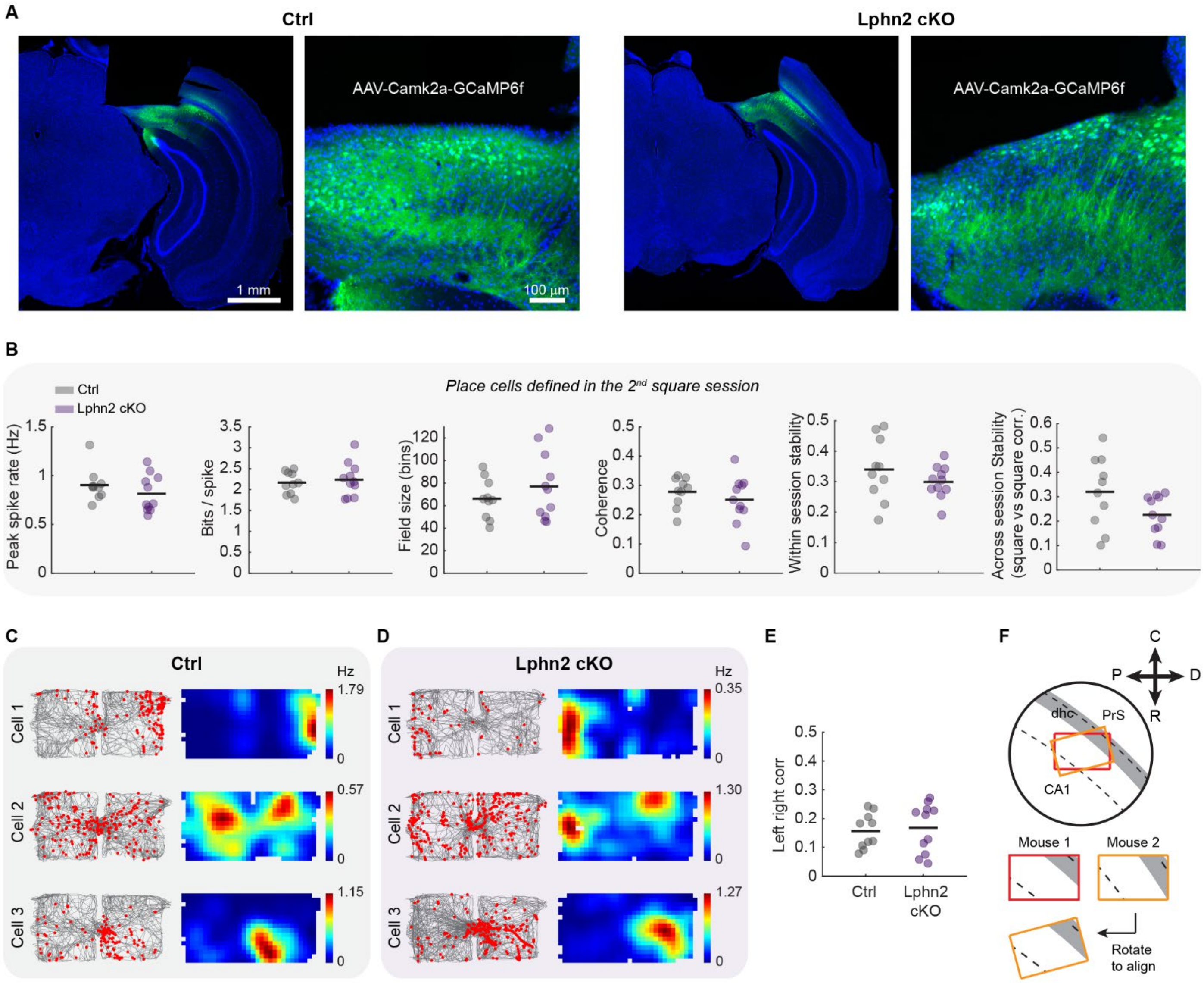


**Figure S1. Histology and tuning property comparisons of subicular place cells, related to Figure 1.**

**(A)** Histology of GRIN lens implantation in the dorsal subiculum of a Ctrl and a *Lphn2* cKO mouse. Green: GCaMP6, Blue: DAPI. Right, enlarged view of GCaMP6-expressing subiculum neurons.

**(B)** Property comparisons between place cells in Ctrl and *Lphn2* cKO mice (Wilcoxon rank-sum test: all $p > 0.05$).

**(C)** Three representative place cells in the shuttle box environment from three Ctrl mice, plotted as in Figure 1D.

**(D)** Same as C, but for *Lphn2* cKO mice.

**(E)** Place cell remapping between the left and right shuttle box compartments (Wilcoxon rank-sum test: $p > 0.05$).

**(F)** Illustration showing miniscope image alignment to ensure a consistent axis across different mice.

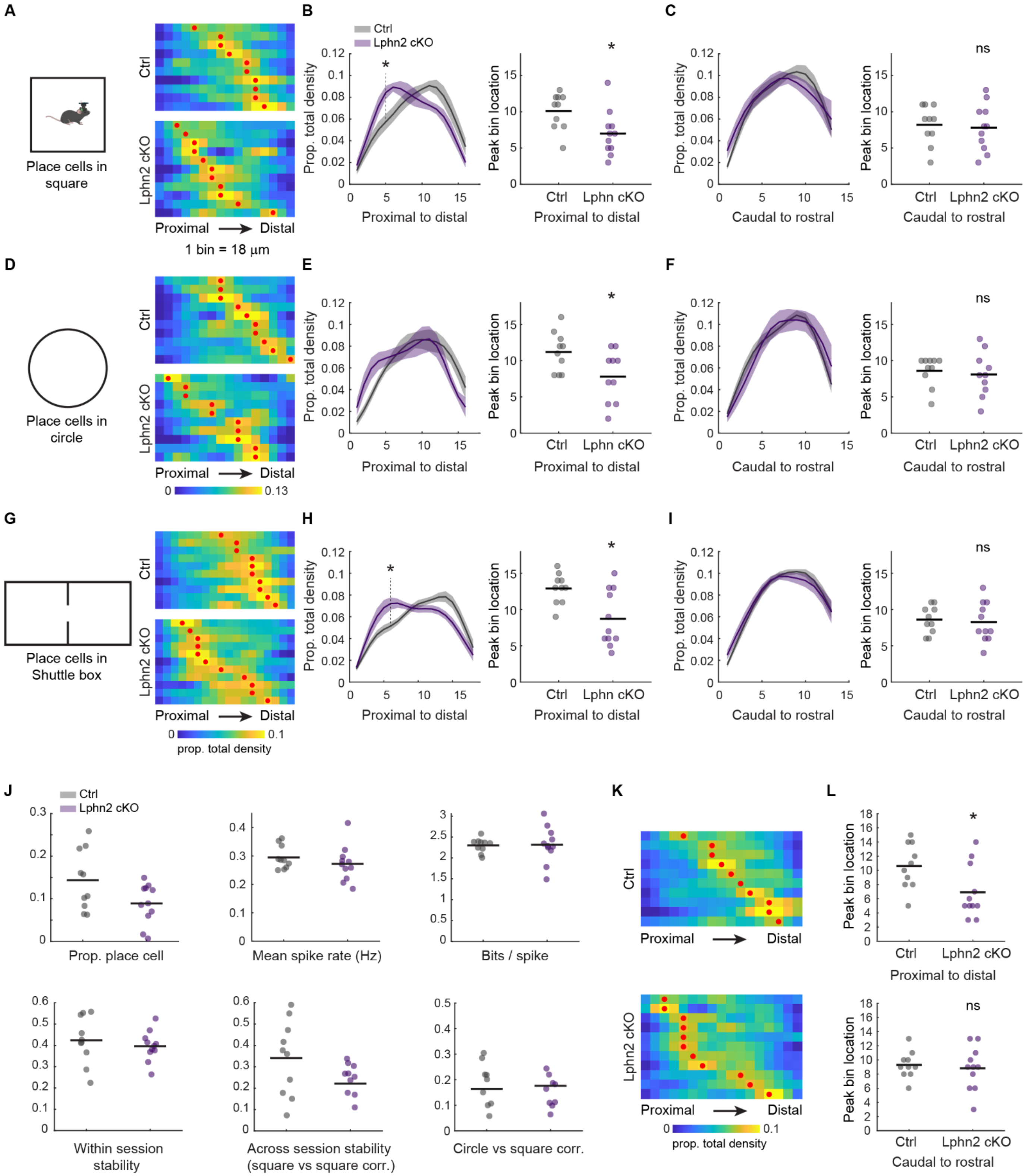


**Figure S2. Comparisons of place cell anatomical distributions within each environment, related to Figure 2.**

**(A)** Distribution of place cell density along the proximodistal axis for all Ctrl (left) and *Lphn2* cKO (right) mice in the square environment. For the heat map, each row represents a mouse, with a red dot indicating the peak of its distribution.

**(B)** Left: Comparison of place cell density distribution along the proximodistal axis between Ctrl and *Lphn2* cKO mice (Ctrl: $n$ = 10 mice, *Lphn2* cKO: $n$ = 11 mice; two-way ANOVA with Sidak's multiple comparisons: * $p$ = 0.046). Right: Peak location of place cell density distributions (Wilcoxon rank-sum test: $p$ = 0.021).

**(C)** Same as B, but along the caudal-to-rostral axis.
**(D)** Same as A, but in the circle environment.
**(E)** Same as B, but in the circle environment (Ctrl: *n* = 10 mice, *Lphn2* cKO: *n* = 10 mice; Right, Wilcoxon rank-sum test: *p* = 0.039).
**(F)** Same as C, but in the circle environment.
**(G)** Same as A, but in the shuttle box environment.
**(H)** Same as B, but in the shuttle box environment (Ctrl: *n* = 10 mice, *Lphn2* cKO: *n* = 11 mice; Left, two-way ANOVA with Sidak's multiple comparisons: * *p* = 0.021; Right, Wilcoxon rank-sum test: *p* = 0.017).
**(I)** Same as C, but in the shuttle box environment.
**(J-L)** Place cells were redefined by adding a coherence criterion (coherence > 0.25). Under this criterion, we compared place cell tuning properties (J) between Ctrl and *Lphn2* cKO mice (Ctrl: n = 10 mice; *Lphn2* cKO: n = 11 mice; Wilcoxon rank-sum test, all *p* > 0.05) and examined their anatomical density distributions (K and L) along the proximodistal and rostrocaudal axes (* *p* = 0.030).

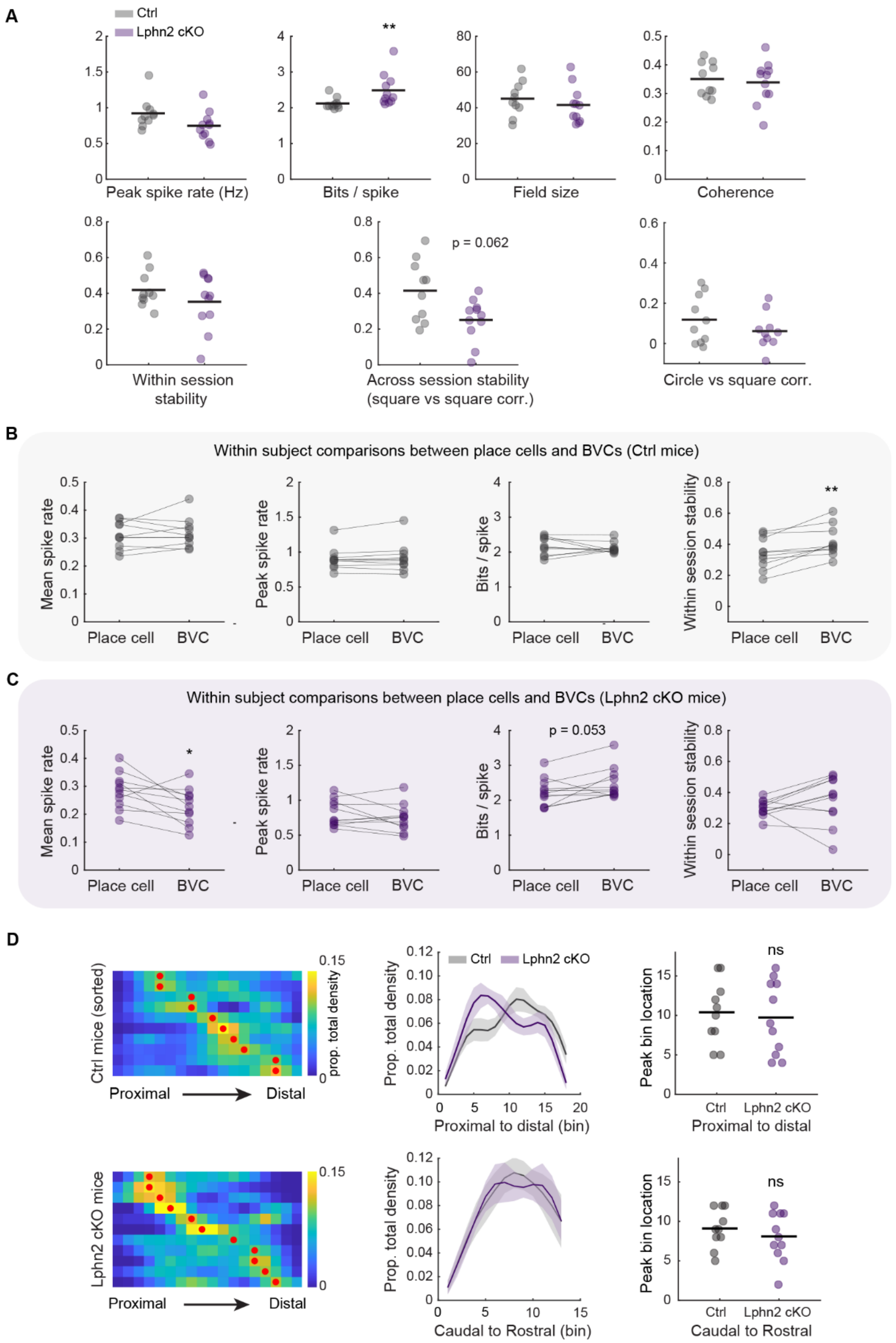


**Figure S3. The effect of *Lphn2* cKO on boundary and corner coding in the subiculum, related to Figure 3.**

**(A)** Tuning property comparisons of boundary vector cells (BVCs) between Ctrl and *Lphn2* cKO mice (Ctrl: $n$ = 10 mice, *Lphn2* cKO: $n$ = 11 mice; Wilcoxon rank-sum test: ** $p$ = 0.0043).

**(B)** Tuning property comparisons between place cells and BVCs within Ctrl mice. (Wilcoxon sign-rank test: ** $p$ = 0.0020).

**(C)** Same as B, but for *Lphn2* cKO mice (Wilcoxon sign-rank test: * $p$ = 0.024).

**(D)** Anatomical distributions and comparisons of corner cell density between Ctrl and *Lphn2* cKO mice, plotted as in Figure 3F–3J.

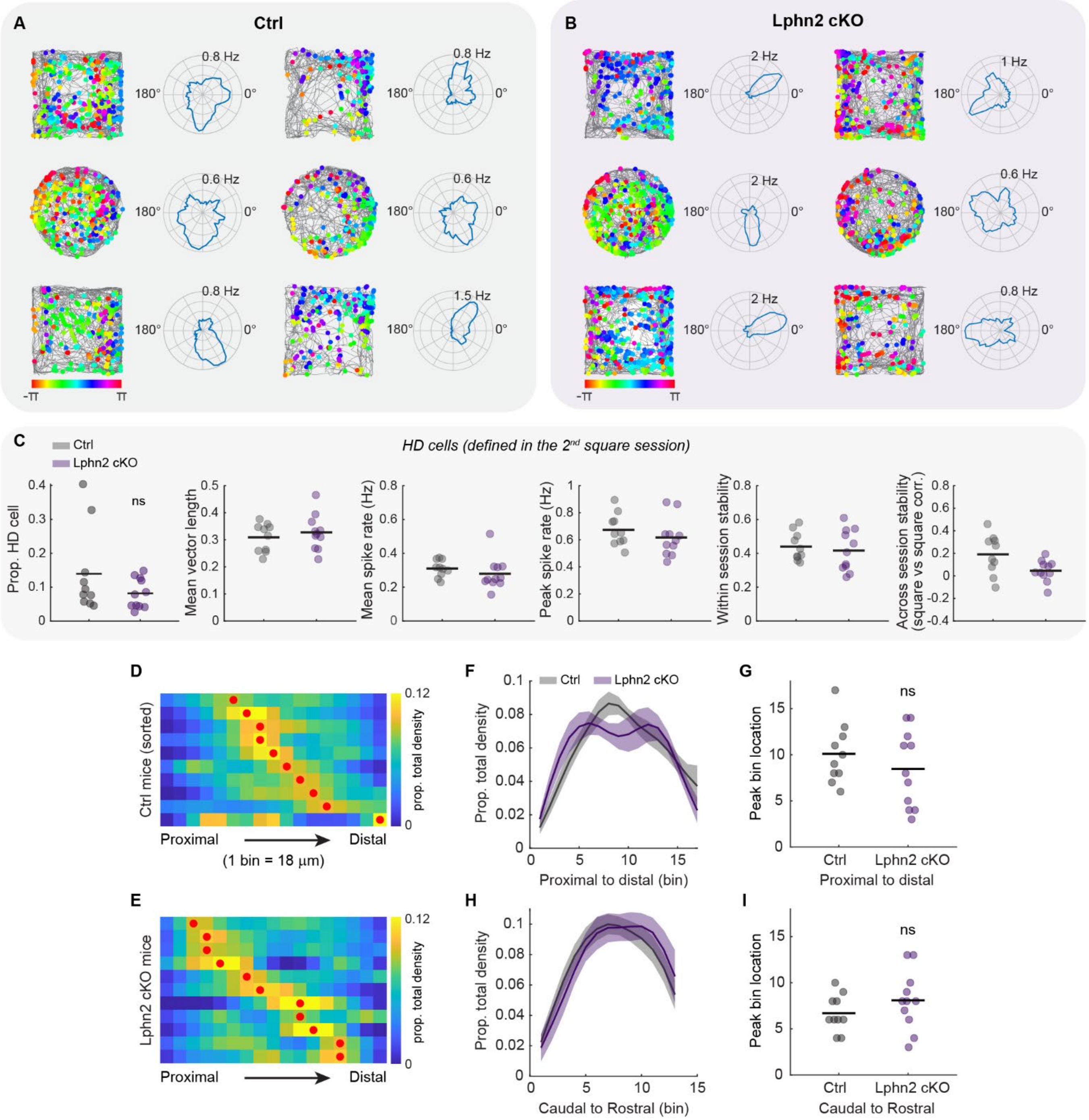


**Figure S4. Head direction tuning in the subiculum remain unaffected in *Lphn2* cKO mice.**

**(A)** Two representative head direction (HD) cells from two Ctrl mice. Each column represents a single HD cell, with its activity tracked across multiple sessions. The raster plot (left) shows deconvolved spikes, which color-coded with animal's allocentric head direction, overlaid on the animal's running trajectory (grey lines). Right shows the polar plots for head direction tuning.

**(B)** Same as A, but for *Lphn2* cKO mice.

**(C)** Tuning property comparisons of HD cells between Ctrl and *Lphn2* cKO mice (Ctrl: $n$ = 10 mice, *Lphn2* cKO: $n$ = 11 mice; Wilcoxon rank-sum test: all $p > 0.05$).

**(D)** Distribution of HD cell density along the proximodistal axis for all Ctrl mice, averaged across square and circle environments. For the heat map, each row represents a mouse, with the red dot indicating the peak of its distribution.

**(E)** Same as D, but for *Lphn2* cKO mice.

**(F)** Comparison of HD cell density distribution along the proximodistal axis between Ctrl and *Lphn2* cKO mice (two-way ANOVA with Sidak's multiple comparisons: all $p > 0.05$).
**(G)** Peak location of place cell density distributions (Wilcoxon rank-sum test: $p = 0.42$).
**(H–I),** Same as F–G, but along the caudal to rostral axis.

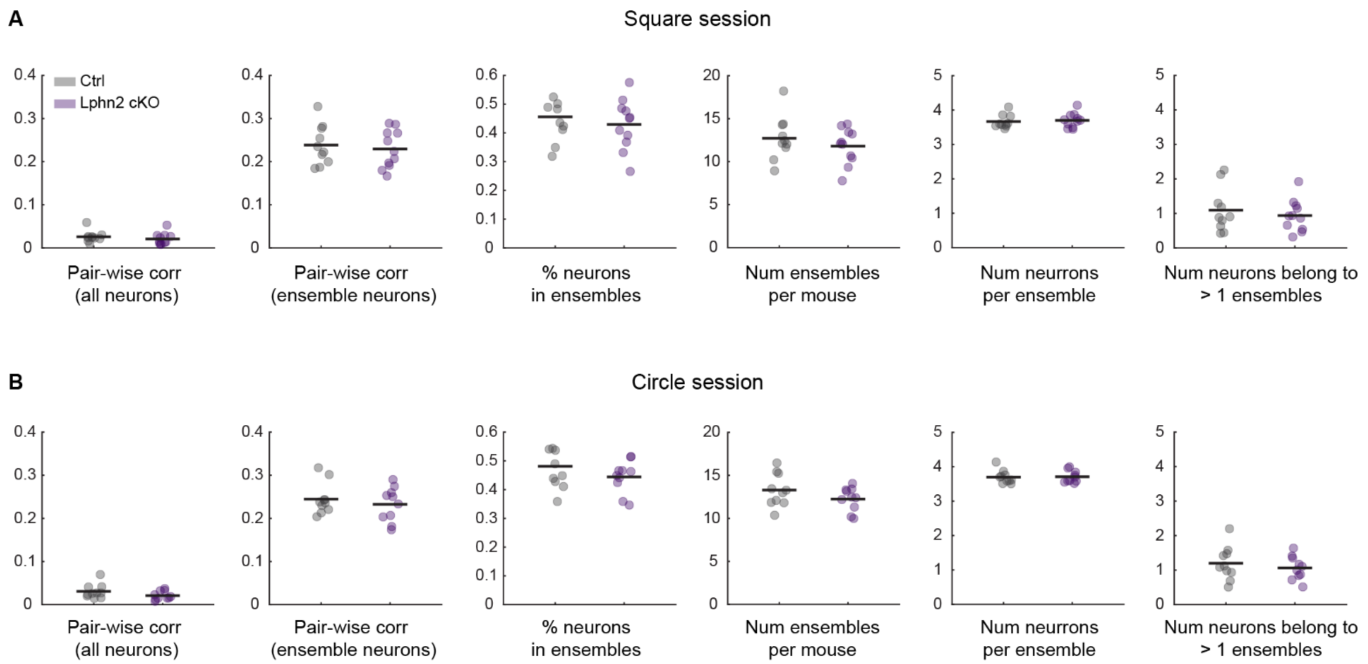


**Figure S5. Property comparisons for coordinated cell ensemble analysis, related to Figure 4.**

**(A)** Comparison of neural ensemble properties (temporal dynamics of calcium signals) between Ctrl ($n$ = 10) and *Lphn2* cKO ($n$ = 11) mice in the square environment. To ensure a fair comparison, the number of neurons in each mouse was first randomly down sampled to 100 for each ensemble detection. This detection process was then run for 250 times, and the final result for each mouse was averaged across the 250 runs (Wilcoxon rank-sum test: all $p > 0.05$).

**(B)** Same as in A, but in the circle environment.
